# Reply to: Eigenmodes of the brain: revisiting connectomics and geometry

**DOI:** 10.1101/2024.08.21.608487

**Authors:** James C. Pang, Kevin M. Aquino, Marianne Oldehinkel, Peter A. Robinson, Ben D. Fulcher, Michael Breakspear, Alex Fornito

## Abstract

In Pang, Aquino et al. (2023)^1^, we presented multiple lines of evidence to indicate that brain geometry plays a previously under-appreciated role in shaping dynamics. Mansour et al. raise concerns about one specific analysis, in which we showed that eigenmodes derived from the geometry of the human cortex can reconstruct diverse activity maps generated with functional magnetic resonance imaging (fMRI) better than eigenmodes derived from connectomes estimated with diffusion MRI (dMRI). Here, we address their concerns and show how our findings and conclusions remain valid.

## Introduction

Mansour et al. motivate their work by claiming that our findings have “been widely interpreted to mean that geometry imposes stronger constraints on cortical dynamics than connectivity”. This interpretation rests on an artificial and inaccurate dichotomy between brain geometry and connectivity, as detailed in the extensive supplementary material (Section S8) of our original article^1^ showing the precise mathematical relation between geometry and connectivity, and our follow-up piece further explaining why the dichotomy is incorrect^2^. To clarify, geometric eigenmodes assume a specific form of connectivity in which cortical locations are coupled through an isotropic, distance-dependent kernel, such that connectivity between any two points decays as an approximately exponential function of their physical separation. This assumption follows the well-known exponential distance rule (EDR) demonstrated empirically to dominate the connectomes of diverse species^3–8^. Geometric eigenmodes thus account for the effects of both cortical geometry and EDR-like connectivity. In contrast, connectome eigenmodes do not directly account for geometry. Connectomes are dominated by EDR-like connectivity^4,9^, but they also account for the effects of topologically complex connections that are not incorporated in an EDR approximation^4,10^.

The stronger performance of geometric eigenmodes in our original analysis^1^ indicated that geometric eigenmodes, and the connectivity approximation that they entail, are sufficient to account for diverse fMRI activity maps, despite the simplicity of the model. We thus concluded that geometric eigenmodes “provide a more compact, accurate, and parsimonious representation of [the brain’s] macroscale activity than alternative connectome-based models” and that “the comparatively poor performance of connectome eigenmodes indicates that topologically complex connections that exist beyond a simple EDR afford minimal further benefit in obtaining eigenmodes that can accurately explain spatiotemporal patterns of cortical activity as measured with fMRI” (p572)^1^.

Mansour et al. focus almost exclusively on our specific comment about the “comparative performance of connectome eigenmodes”, citing it multiple times. They report a thorough set of analyses to address this “comparatively poor performance” by showing that, when dMRI data are processed using a specific pipeline of “state-of-the-art connectome reconstruction techniques”, the accuracy of the connectome and geometric models becomes approximately equal. We applaud their efforts to identify a dMRI processing regime that improves the accuracy of the connectome model. However, we disagree with their conclusion that the “evidence presented to support the comparative proposition that “eigenmodes derived from brain geometry represent a more fundamental anatomical constraint on dynamics than the connectome” may require reconsideration.” Below, we explain why their findings only strengthen our original claims.

### Geometric eigenmodes are more parsimonious than connectome eigenmodes

Mansour et al. consider five dMRI processing steps that can mitigate “biases and inaccuracies” associated with “high-resolution connectome mapping”. Each of these steps requires investigator-dependent choices between alternative approaches that substantially influence connectivity estimates and model accuracy, as thoroughly detailed in Mansour et al.’s 10 multi-panel supplementary figures (their Figs. S1–10). For instance, their two different methods for gyral bias correction yield connectomes with quite different architectures (Figs. 1–2 and their Fig. S1), neither of which completely address the gyral bias that was suggested to contaminate our original connectome (Fig. 2b). This simple methodological variation is sufficient to reduce the correlation between the two resulting connectivity matrices to 0.46, despite the source data being otherwise identical. This major discrepancy in connectivity weights, caused by a single processing choice, underscores the fragility of the resulting connectomes (as suggested in our own original analysis^1^) and poses a difficult challenge for the construction of a reliable basis set, particularly given the priority that Mansour et al. assign to the analysis of weighted connectomes.

**Figure 1.**
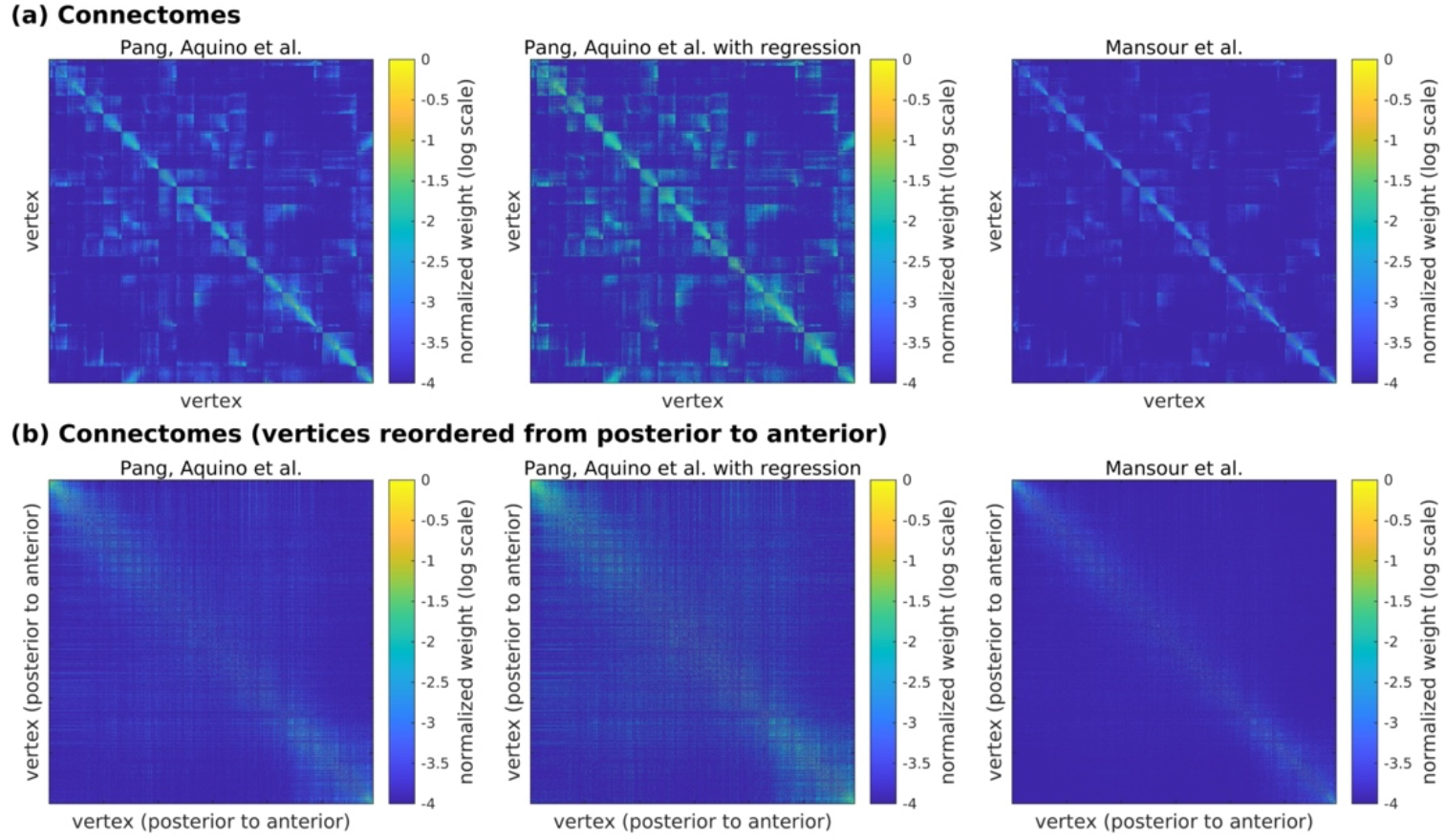
Original Pang, Aquino et al. and reprocessed Mansour et al. connectomes. (**a**) Panels from left to right show the original connectome used in our work, our connectome after performing Mansour et al.’s gyral bias correction via regression, and the connectome preferred by Mansour et al. (**b**) Same as panel a but the vertices are reordered according to their spatial locations from posterior to anterior. This reordering shows how the connectome preferred by Mansour et al. attenuates long-range connectivity, thus converging on the distance-dependent connectivity approximation inherent in the geometric approach.

**Figure 2.**
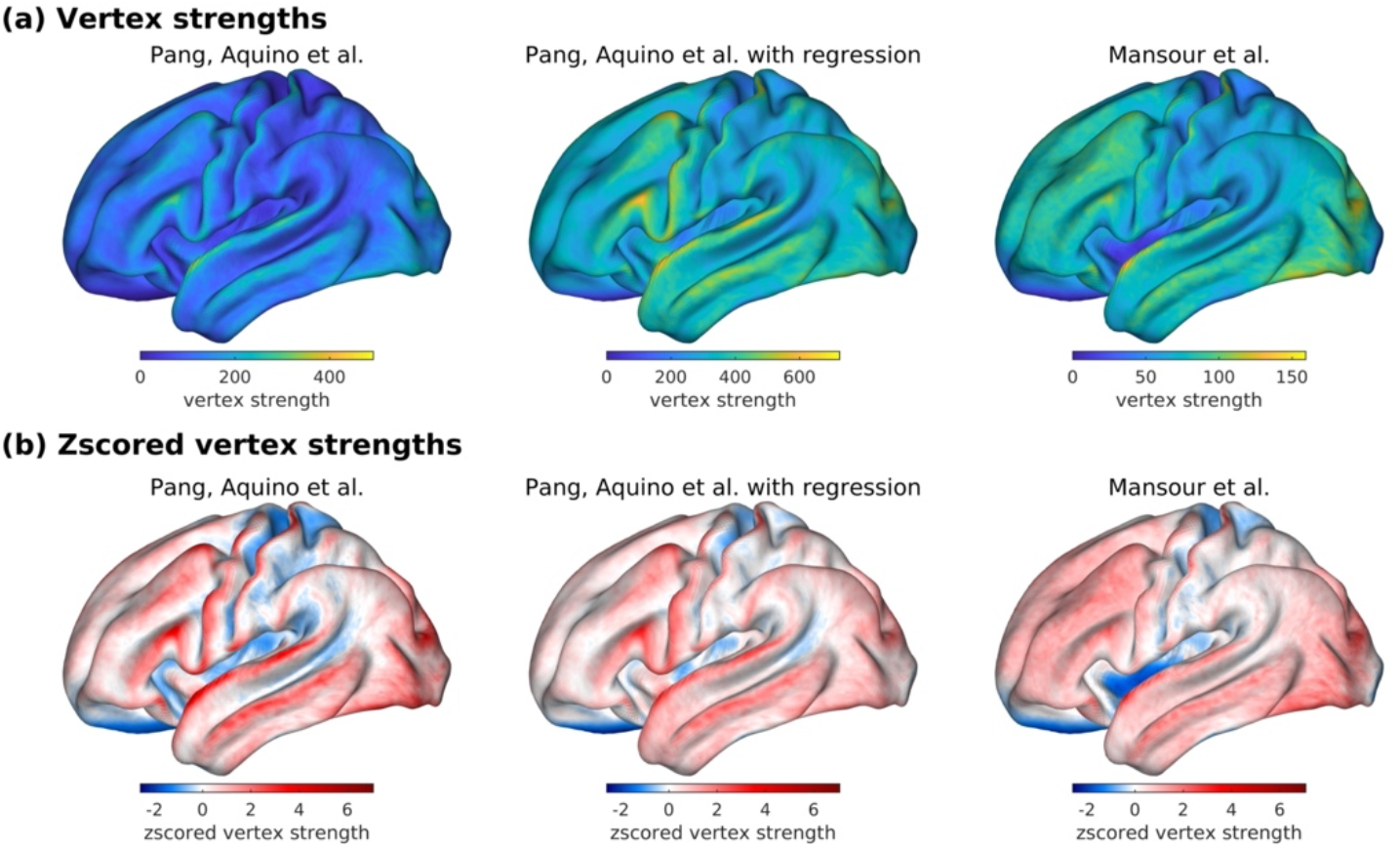
Spatial maps of vertex-level connectivity strengths for the original and reprocessed connectomes. (**a**) Spatial map of the vertex connectivity strengths as depicted by Mansour et al. (**b**) Same as panel a but the values are zscored, which attenuates the effect of outlying values and allows a clearer visualization of the spatial structure in the map. For panels a and b, panels from left to right show results using our original connectome, our connectome after performing Mansour et al.’s gyral bias correction via regression, and Mansour et al.’s preferred connectome. Panel b clearly shows that gyral bias is still evident in the reprocessed connectomes and that the two correction strategies by Mansour et al. can lead to highly divergent maps. These visualizations are preferable to the regression slopes and linear correlations shown in Fig. S1b of Mansour et al., which will be biased by the outliers, heteroscedasticity, and non-monotonicity evident in their scatterplots of the association between vertex connectivity strength and curvature.

Notably, this critical choice of gyral bias correction is only one in a sequence of decisions required in their connectome reconstruction pipeline. These decisions yield a parameter space comprising many plausible processing steps that can result in very different connectomes^11,12^. It is difficult to ascertain how one should choose between these different processing options. Mansour et al. use the geometric eigenmodes as a benchmark for optimizing their set of choices. This approach is vulnerable to overfitting the data to obtain a desired result. It is unclear how well this optimized set of choices generalizes to independent data.

Geometric eigenmodes obviate the need to navigate this analytic “garden of forking paths”. They are derived using robust and widely accepted pipelines for reconstructing cortical surfaces from T1-weighted scans^13^. In fact, the end point of the geometric eigenmode pipeline is the starting point of the connectome eigenmode pipeline, which requires acquisition of an additional MRI modality and the construction of a complex sequence of processing steps based on numerous choices. Mansour et al.’s analysis shows that, at best, a specific subset of these choices yields a basis set that matches, but never substantially surpasses, the accuracy of the geometric eigenmodes (their Figs 1, S2, S4–10, and S12). The fact that equivalent performance is the best result for the connectome approach, despite the large parameter space explored by Mansour et al., is strong evidence for the superior parsimony of the geometric basis set. Their analysis thus strengthens our conclusion that geometric eigenmodes offer a sufficient and parsimonious account of fMRI data. Future innovations in dMRI processing may further improve the performance of connectome eigenmodes, but the gain in model performance should be of a sufficient magnitude to justify the added complexity and problematic reliability of the connectome model.

### Topologically complex connections make a negligible contribution to reconstruction accuracy

The ability to account for the specificity of brain connectivity, and particularly of the topologically complex, often long-range, connections that cannot be explained by an EDR, is the primary motivation for using connectome eigenmodes. However, Mansour et al.’s own findings show that removal of longer-range connections (i.e., >16 mm) has a minimal effect on the reconstruction accuracy of connectome eigenmodes, whereas removal of short-range connections has a much larger impact (their Fig. S12). This result, when taken with their finding of approximately equivalent performance of the geometric and “state-of-the art” connectome eigenmodes, strengthens our original claim that an isotropic, EDR-like connectivity approximation is sufficient to reconstruct diverse activity patterns mapped with fMRI and that topologically complex connectivity (i.e., connectivity not readily captured by an EDR-like approximation) makes a minimal contribution. Mansour et al.’s demonstration that removal of short-range connections has a larger impact on reconstruction accuracy also supports our original claim that local, short-range connections, which facilitate continuous propagation of waves of excitation through the cortex, are important for understanding fMRI activity^1^. As we acknowledged in our original article^1^, topologically complex connections undoubtedly play important roles for brain function but their effects may not be easily revealed by traditional fMRI paradigms.

### “State-of-the-art” dMRI processing increases the similarity between connectome and geometric eigenmodes

If long-range connections make a negligible contribution to model accuracy, what drives the performance improvement of “state-of-the-art” connectome eigenmodes? The answer to this question is found in Figs 2 and S11b of Mansour et al., which show that their processing strategy increases the alignment of the connectome eigenmodes to the geometric eigenmodes. This increased similarity occurs because their additional processing steps accentuate the local homogeneity and EDR-like dependence of the connectome––the two key elements of the connectivity approximation upon which the geometric approach relies. For instance, their Figs S3f–g show how local homogeneity and the EDR dependence are dramatically enhanced by connectome spatial smoothing. In Fig. 3a, we additionally demonstrate that their gyral bias correction truncates the tails of the vertex connectivity strength distribution relative to the uncorrected case, yielding an approximately Gaussian shape in which there is a low probability of finding vertices with substantially higher or lower connectivity than the mean. Thus, regardless of any debate over whether gyral bias correction improves the biological plausibility of vertex-wise connectivity estimates, it ultimately increases the homogeneity of those estimates. Figure 3b also shows that the processing steps applied by Mansour et al. exaggerate the distance-dependent decay of connectivity edge weights by attenuating precisely those longer-range connections that deviate from an EDR footprint (see also Fig. 1b).

**Figure 3.**
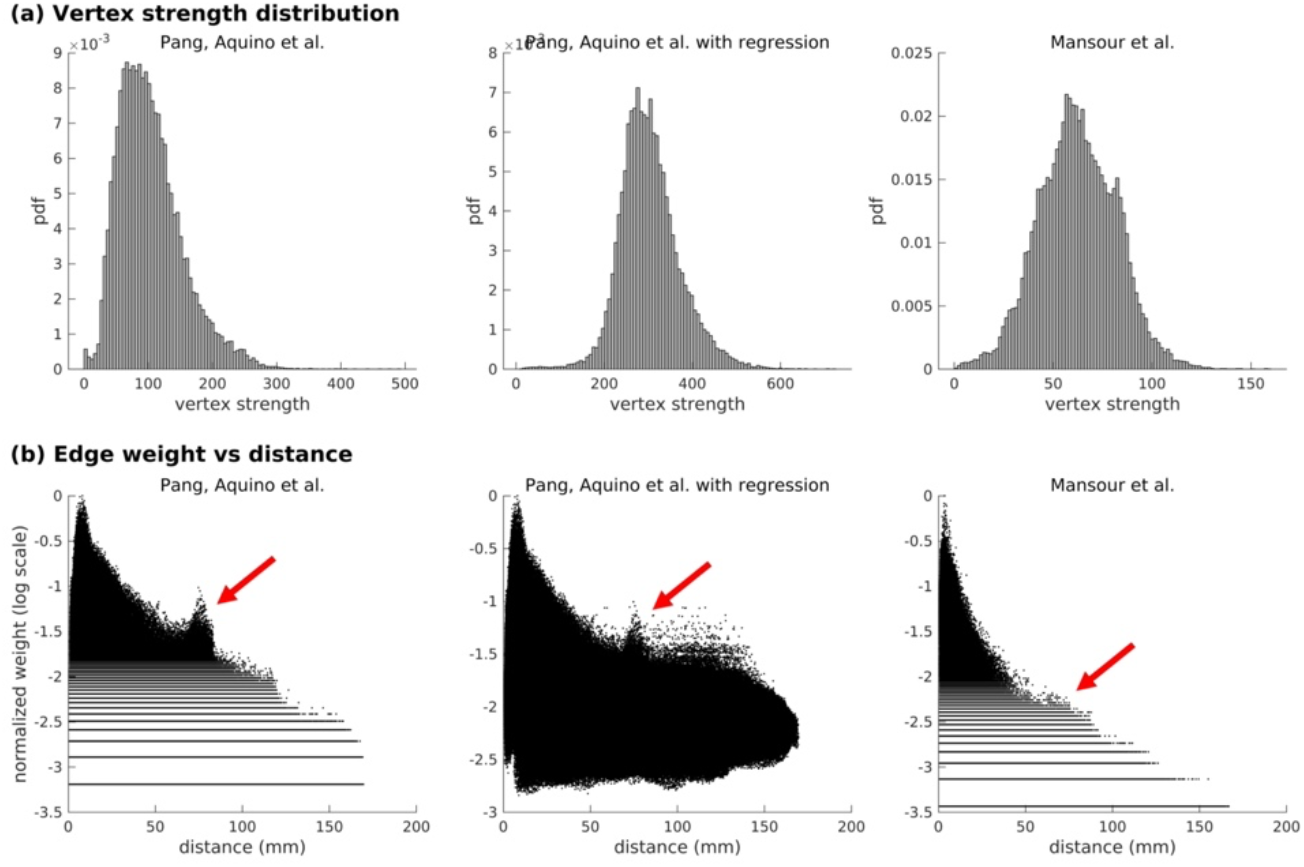
Vertex connectivity strength distributions and edge weight distance-dependencies for the original and reprocessed connectomes. (**a**) Distribution of vertex connectivity strengths. (**b**) Edge weight vs euclidean distance between vertices in semilogarithmic scale. For panels a and b, left to right shows results using our original connectome, our connectome after performing Mansour et al.’s gyral bias correction via regression, and Mansour et al.’s preferred connectome. The processing steps proposed by Mansour et al. truncate the tails of the vertex connectivity strength distribution (panel a) and attenuate the weights of medium-to-long-range connections (between 50 and 100 mm) (panel b; see red arrows). The processing steps of Mansour et al. thus enforce a structure on the empirical data that more closely resembles the locally isotropic, EDR-like connectivity approximation of the geometric model.

These considerations indicate that the performance of connectome eigenmodes approaches geometric eigenmodes when the former are made to look like the latter. This is done through data processing steps that increase the convergence between empirical connectivity estimates and the isotropic, EDR-like connectivity approximation that underpins the geometric model. The findings of Mansour et al. thus align with our original results and support the fundamental role of geometric eigenmodes and the connectivity approximation that they entail in shaping fMRI activity maps.

## Conclusions

Mansour et al. show that: (i) best-estimate connectome eigenmodes only ever match, but never substantially surpass, the reconstruction accuracy of the geometric model; (ii) long-range connections make a minimal contribution to the performance of the connectome model; and (iii) the performance improvements of the connectome model are attributable to processing steps that increase the similarity between empirical connectomes and the connectivity approximation inherent in the geometric approach. These findings beg the question: why use the more complex connectome model if it offers little added value beyond the simpler geometric approach? The answer to this question depends on the goals of the investigator, since both geometric and connectome eigenmodes have distinct strengths and limitations. Geometric eigenmodes offer a simple and parsimonious account of brain function as they are formally linked with the biophysical model of dynamics provided by neural field theory^14–17^. This link allows one to associate each eigenmode with characteristic spatial and temporal frequencies, which can be used to understand how different types of stimuli drive cortical dynamics^18^. However, a limitation of the geometric approach is that it is not straightforward to integrate cortical and subcortical eigenmodes into a single model. Connectome eigenmodes can account for both cortical and subcortical connectivity and facilitate mappings between specific eigenmodes and task activation patterns in specific circumstances (see Fig. S13 of Mansour et al.). However, connectome eigenmodes are susceptible to myriad data processing choices and include overlapping contributions from topologically complex connections, EDR-like connectivity, and geometry that can be difficult to disentangle without additional analysis. Future work will benefit from understanding the strengths and limitations of the two approaches, and developing approaches to unify them.

## References

1. Pang, J. C. et al. Geometric constraints on human brain function. Nature 618, 566–574 (2023).

2. Pang, J. C. et al. Reply to: Commentary on Pang et al. (2023) Nature. 2023.10.06.560797 Preprint at 10.1101/2023.10.06.560797 (2023).

3. Henderson, J. A. & Robinson, P. A. Relations between the geometry of cortical gyrification and white-matter network architecture. Brain Connectivity 4, 112–130 (2014).

4. Roberts, J. A. et al. The contribution of geometry to the human connectome. NeuroImage 124, 379–393 (2016).

5. Wang, X. J. & Kennedy, H. Brain structure and dynamics across scales: In search of rules. Current Opinion in Neurobiology 37, 92–98 (2016).

6. Horvát, S. et al. Spatial Embedding and Wiring Cost Constrain the Functional Layout of the Cortical Network of Rodents and Primates. PLOS Biology 14, e1002512 (2016).

7. Ercsey-Ravasz, M. et al. A Predictive Network Model of Cerebral Cortical Connectivity Based on a Distance Rule. Neuron 80, 184–197 (2013).

8. Knoblauch, K., Ercsey-Ravasz, M., Kennedy, H. & Toroczkai, Z. The Brain in Space. in Micro-, Meso- and Macro-Connectomics of the Brain (eds. Kennedy, H., Van Essen, D. C. & Christen, Y.) 45–74 (Springer International Publishing, Cham, 2016). doi:10.1007/978-3-319-27777-6_5.

9. Theodoni, P. et al. Structural Attributes and Principles of the Neocortical Connectome in the Marmoset Monkey. Cerebral Cortex 32, 15–28 (2022).

10. Betzel, R. F. et al. Generative models of the human connectome. NeuroImage 124, 1054–1064 (2016).

11. Oldham, S. et al. The efficacy of different preprocessing steps in reducing motion-related confounds in diffusion MRI connectomics. NeuroImage 222, 117252 (2020).

12. Gajwani, M. et al. Can hubs of the human connectome be identified consistently with diffusion MRI? Network Neuroscience 1–25 (2023) doi:10.1162/netn_a_00324.

13. Fischl, B. FreeSurfer. NeuroImage 62, 774–781 (2012).

14. Robinson, P. A., Rennie, C. J. & Wright, J. J. Propagation and stability of waves of electrical activity in the cerebral cortex. Physical Review E 56, 826–840 (1997).

15. Robinson, P. A. et al. Prediction of electroencephalographic spectra from neurophysiology. Physical Review E 63, 021903 (2001).

16. Jirsa, V. & Haken, H. Field Theory of Electromagnetic Brain Activity. Physical Review Letters 77, 960–963 (1996).

17. Wright, J. J. & Liley, D. T. J. Simulation of electrocortical waves. Biological Cybernetics 72, 347–356 (1995).

18. Gabay, N. C., Babaie-Janvier, T. & Robinson, P. A. Dynamics of cortical activity eigenmodes including standing, traveling, and rotating waves. Phys. Rev. E 98, 042413 (2018).

